# A regulatory interface on RIPK2 is required for XIAP binding and NOD signaling activity

**DOI:** 10.1101/2020.03.12.988725

**Authors:** Valentin J. Heim, Laura F. Dagley, Che A. Stafford, Fynn M. Hansen, Elise Clayer, Aleksandra Bankovacki, Andrew I. Webb, Isabelle S. Lucet, John Silke, Ueli Nachbur

## Abstract

Signaling via the intracellular pathogen receptors Nucleotide-binding oligomerization domain-containing proteins NOD1 and NOD2 requires Receptor Interacting Kinase 2 (RIPK2), an adaptor kinase that can be targeted for the treatment of various inflammatory diseases. However, the molecular mechanisms of how RIPK2 contributes to NOD signaling are not completely understood. We generated FLAG-tagged RIPK2 knock-in mice using CRISPR/Cas9 technology to study NOD signaling mechanisms at the endogenous level. Using cells from these mice we were able to generate a detailed map of post-translational modifications on RIPK2 during NOD signaling and we identified a new regulatory interface on RIPK2, which dictates the crucial interaction with the E3 ligase XIAP.

## Introduction

Nucleotide-binding oligomerization domain-containing (NOD) proteins NOD1 and NOD2 are intracellular pathogen recognition receptors (PRRs) that sense the bacterial peptidoglycan (PGN) fragments *γ*-D-Glu-m diaminopimelic acid (DAP) and muramyl dipeptide (MDP), respectively [1, 2]. NOD1 and NOD2 play an important role in the clearance of bacterial pathogens, including *Mycobacterium tuberculosis* [3], *Listeria monocyt*ogenes [4] and multiple *Chlamydiae* species [5]. Aberrant NOD signaling has long been associated with a range of inflammatory disorders [6, 7], and recent findings suggest that inhibition of the NOD pathways could be beneficial in the treatment of allergic asthma [8], and type 2 diabetes mellitus (T2DM) [9-12].

Binding of the respective ligands to NOD1 and NOD2 leads to their self-oligomerization [13] and the recruitment of Receptor-interacting serine/threonine-protein kinase 2 (RIPK2) via homotypic caspase recruitment domain (CARD)-CARD interactions [14]. RIPK2 is the essential adaptor kinase in the NOD signaling pathway and drives nuclear factor kappa-light-chain-enhancer of activated B cells (NF-*κ*B) and mitogen-activated protein (MAP) kinase activation [15, 16]. The kinase activity of RIPK2 was initially reported to be required for signal transduction and for critical autophosphorylation of RIPK2 on S176 in the activation loop of the kinase domain [17] and Y474 in its CARD [18]. However, recent studies suggest that RIPK2 kinase activity is dispensable for NF-*κ*B activation and cytokine production [19, 20]. Furthermore, it has been established that NOD signaling relies on ubiquitination of RIPK2 [21]. This process is coordinated by multiple ubiquitin E3 ligases, including X-linked inhibitor of apoptosis protein (XIAP) [22-24]. XIAP binds to the kinase domain of RIPK2 via its baculovirus IAP repeat 2 (BIR2) domain [22] [25, 26] and generates K63-linked polyubiquitin chains on multiple lysine residues [19]. This leads to the recruitment of the linear ubiquitin chain assembly complex (LUBAC) [24] and the generation of M1-linked polyubiquitin chains on RIPK2 that serve as binding platforms for I*κ*B kinase (IKK) and Transforming growth factor beta-activated kinase 1 (TAK1) complexes.

While the importance of XIAP and LUBAC for immune responses mediated by NOD1 and NOD2 has been demonstrated *in vitro* and *in vivo*, not much is known about the function of individual ubiquitination sites on RIPK2. A putatively ubiquitinated lysine residue on RIPK2 (K209) was discovered more than 10 years ago in a systematic screening of lysine to arginine mutations (K/R) that disrupted NF-*κ*B activation in overexpression experiments [27]. A subsequent study showed that the K209R mutation blocked RIPK2 ubiquitination and signaling [21], and it was concluded that K209 is directly ubiquitinated and is indispensable for NOD2 responses. Nevertheless, ubiquitination of RIPK2 on K209 has not been shown. Intriguingly, a recently published proteomics experiment reported multiple ubiquitination sites on RIPK2, but the authors did not identify K209 [19]. Instead, they identified ubiquitination sites on the C terminus of RIPK2 and a K410R/K538R double mutation reduced MDP-dependent responses of THP-1 cells. Altogether, this highlights that our understanding of how post-translational modifications (PTMs) of RIPK2 regulate NOD signaling is incomplete.

Due to the association with inflammatory diseases, pharmaceutical companies have invested in the development of inhibitors for the NOD signaling pathway. RIPK2 has been established as a potential drug target, particularly in inflammatory bowel disease, and RIPK2-targeting kinase inhibitors have been developed [24, 26, 28]. Recent studies showed that the inhibition of NOD signaling is not directly due to the inhibition of the kinase function of RIPK2, but rather by the disruption of the RIPK2-XIAP interaction [19, 20, 26]. This has led to the hypothesis that protein-protein interaction inhibitors could be used to treat NOD-driven diseases and highlights the need for a detailed understanding of posttranslational modifications on RIPK2.

A significant issue that has hindered our understanding of such mechanisms of the NOD signaling pathway is the lack of specific biochemical tools. Most studies have been limited to overexpression experiments in cancer cell lines, which has many drawbacks including the formation of artefactual interactions or altered protein activities [29, 30]. In the context of NOD signaling, it was shown that ectopic overexpression of NOD receptor complex components leads to pathway activation independent of PGN binding [31-33]. To study NOD2 signaling mechanisms and investigate the molecular determinants of RIPK2 activation, we established a new mouse strain with endogenously FLAG-tagged RIPK2. This allowed us to isolate RIPK2 from primary tissues and cells and to characterize the PTMs on RIPK2 that occur during MDP-induced signaling at endogenous levels. Using this approach, we identified a novel regulatory interface that mediates XIAP binding and is required for signal transduction.

## Results

### FLAG-RIPK2 knock-in mice represent a novel tool to study endogenous NOD signaling mechanisms

To study NOD signaling at the endogenous level and to investigate how RIPK2 regulates signal transduction, we generated N-terminally FLAG tagged RIPK2 knock-in mice by microinjection of single guide RNAs (sgRNAs), recombinant Cas9 protein and a dsDNA oligonucleotide encoding the FLAG-tagged sequence of RIPK2 with homologous arms upstream and downstream of the sgRNA targeted region into wild-type C57Bl/6 embryos. Using this process, we generated mice harboring the desired FLAG-tagged version of RIPK2, as well as mice with defined insertions and deletions. After backcrossing to C57Bl/6 mice, we established a FLAG-RIPK2 knock-in mouse strain as well as a new RIPK2 knock-out strain.

First, we explored the tissue distribution of RIPK2 by testing homogenates from organs of FLAG-RIPK2 and wild-type mice by Western Blot. As the expression levels of RIPK2 in these organs were too low for detection using anti-FLAG™ antibodies, we therefore subjected organ homogenates to anti-FLAG immunoprecipitation (**Fig. 1A**) and probed the supernatants from boiled beads for the presence of RIPK2. We found that FLAG-RIPK2 was readily expressed in the brain, colon, small intestine, lung, skin, liver and spleen but we did not detect FLAG-RIPK2 in the kidney.

**Figure 1.**
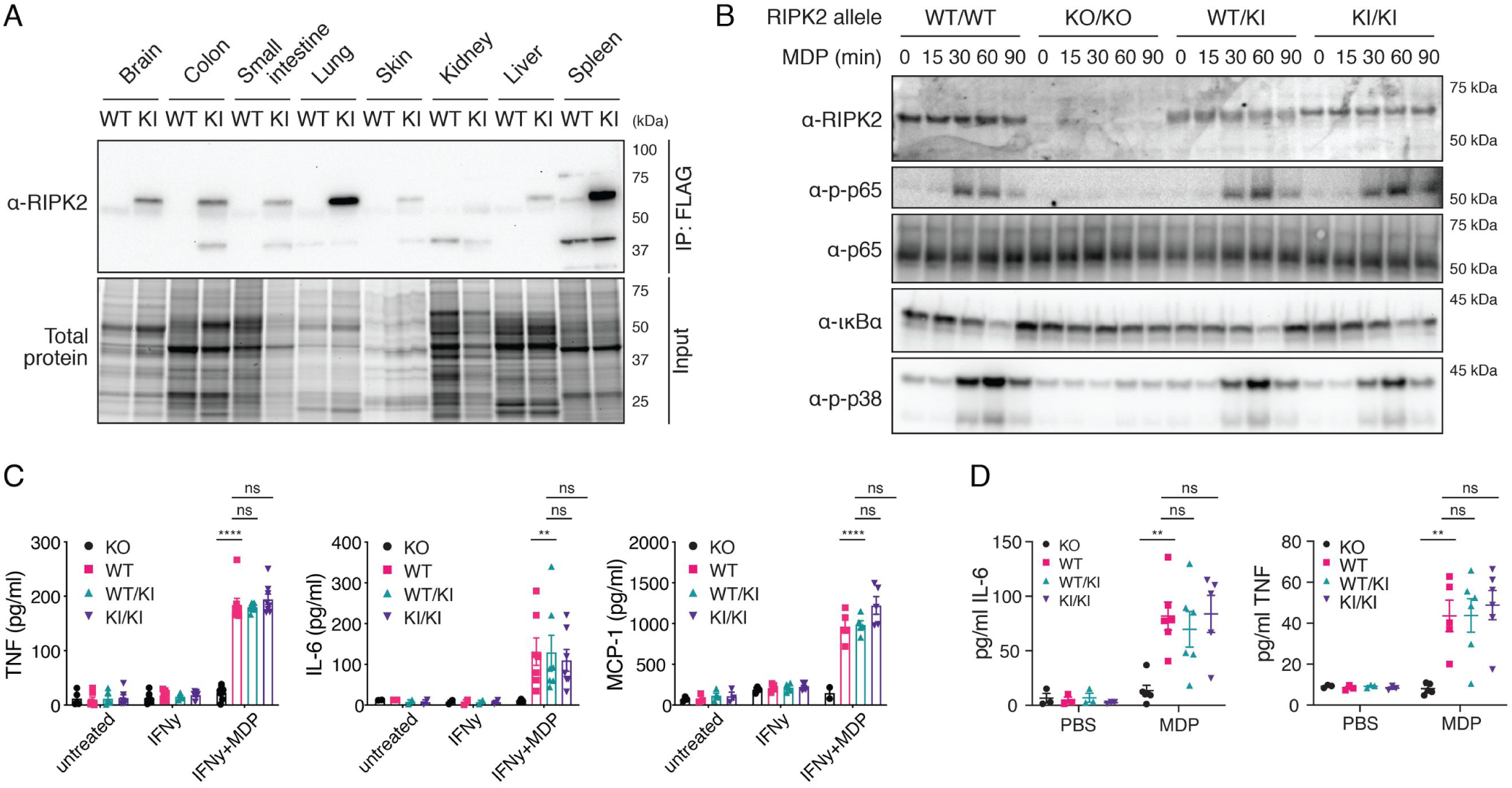
Flag-RIPK2 knock-in mice as a tool to study endogenous NOD signaling mechanisms. A Tissue distribution of RIPK2 determined by anti-FLAG immunoprecipitation. Organ homogenates from WT (wild-type) and KI (homozygous FLAG-RIPK2) mice were subjected to anti-FLAG immunoprecipitation and immunoblotting. B Inflammatory signaling in wild-type (WT/WT), RIPK2 CRISPR KO (KO/KO) and FLAG-RIPK2 heterozygous (WT/KI) and homozygous (KI/KI) BMDMs. BMDMs were primed with IFNy, stimulated with MDP for indicated times and analysed by immunoblotting. C Cytokine production of RIPK2 CRISPR KO (KO), wild-type (WT) and FLAG-RIPK2 heterozygous (WT/KI) and homozygous (KI/KI) BMDMs in response to MDP. BMDMs were left untreated or treated with IFNy or IFNy and MDP over-night and cytokines were measured by ELISA. N = 5-8 mice. Shown is average ± SEM. ns = P > 0.05; * = P ≤ 0.05; ** = P ≤ 0.01; *** = P ≤ 0.001; **** = P ≤ 0.0001. D Serum cytokines in RIPK2 CRISPR KO (KO), wild-type (WT) and FLAG-RIPK2 heterozygous (WT/KI) or homozygous (KI/KI) mice after i.p. MDP administration. Mice were injected i.p. with PBS or MDP, sacrificed after 4h and serum cytokines were measured by ELISA. N = 3-6 mice. Shown is average ± SEM. ns = P > 0.05; * = P ≤ 0.05; ** = P ≤ 0.01; *** = P ≤ 0.001; **** = P ≤ 0.0001.

We tested the functionality of FLAG-RIPK2 using IFN*γ*-primed bone marrow-derived macrophages (BMDMs). Upon MDP stimulation, cells generated from homozygous (KI/KI) or heterozygous (WT/KI) FLAG-RIPK2 mice induced NF-*κ*B and MAP kinase pathways equivalent to wild-type cells (**Fig. 1B**). Western Blot for RIPK2 also revealed even expression levels between wild-type and FLAG-tagged versions of RIPK2. As expected, cells generated from our new strain of RIPK2 knock-out mice were unresponsive to MDP and did not express detectable RIPK2. Cytokine production of BMDMs after MDP stimulation was then measured by ELISA (**Fig. 1C**). In IFN*γ*-primed wild-type and FLAG-RIPK2 BMDMs, treatment of MDP induced the secretion of TNF, IL-6 and MCP-1 at equal levels, while RIPK2 KO BMDMs were unresponsive. To confirm that NOD2-dependent responses in FLAG-RIPK2 mice were indistinguishable from wild-type mice *in vivo*, we intraperitoneally (i.p.) injected MDP or PBS into wild-type, FLAG-RIPK2 and RIPK2 knock-out mice and measured cytokine levels in the serum by ELISA (**Fig. 1D**). MDP challenge caused a reproducible increase in the levels of Il-6 and TNF in the serum of wild-type and knock-in mice. In RIPK2 deficient mice, MDP injection did not result in an increase of cytokines. Altogether these data show that FLAG-RIPK2 mice and primary cells from these mice responded normally to NOD2 stimulation.

### Post-translational modifications on RIPK2

As described above, phosphorylation and ubiquitination of RIPK2 are critical for the assembly of the NOD2 signaling complex and downstream signaling. Despite the known modifications that RIPK2 undergoes during signaling we rarely observe ubiquitination of RIPK2, characterized by a high molecular weight smear in Western blots, in whole-cell lysates of MDP-stimulated cells. Accordingly, we did not observe PTMs on RIPK2 when we performed an anti-FLAG immunoprecipitation followed by mass spectrometry analysis on MDP-stimulated FLAG-RIPK2 BMDMs. This indicates that the majority of cellular RIPK2 is not part of the NOD signaling complex, even after stimulation, and suggests that an additional purification step to investigate RIPK2 in its activated state is required.

To enrich for the RIPK2 pool that participated in NOD2 signaling, we established a sequential pulldown protocol. IFN*γ*-primed BMDMs were stimulated with MDP for 30 minutes and lysates were first enriched for ubiquitinated proteins using glutathione S-transferase (GST)-ubiquitin associated domain (UBA) bound to Sepharose beads [34]. Bound proteins were then eluted with Glutathione, and eluates were subjected to anti-FLAG immunoprecipitation, followed by elution with 3x-FLAG peptide (**Fig. 2A**). The first pulldown (UBA, lane B) yielded a sample containing readily detectable levels of a ladder of RIPK2 species suggestive of ubiquitination, as well as many other ubiquitinated proteins (anti-ubiquitin, bottom panel). The sample obtained by sequential pulldown with anti-FLAG (lane C) also contained modified RIPK2 however the background of ubiquitinated proteins was much reduced, further suggesting that this approach resulted in purification of activated RIPK2.

**Figure 2:**
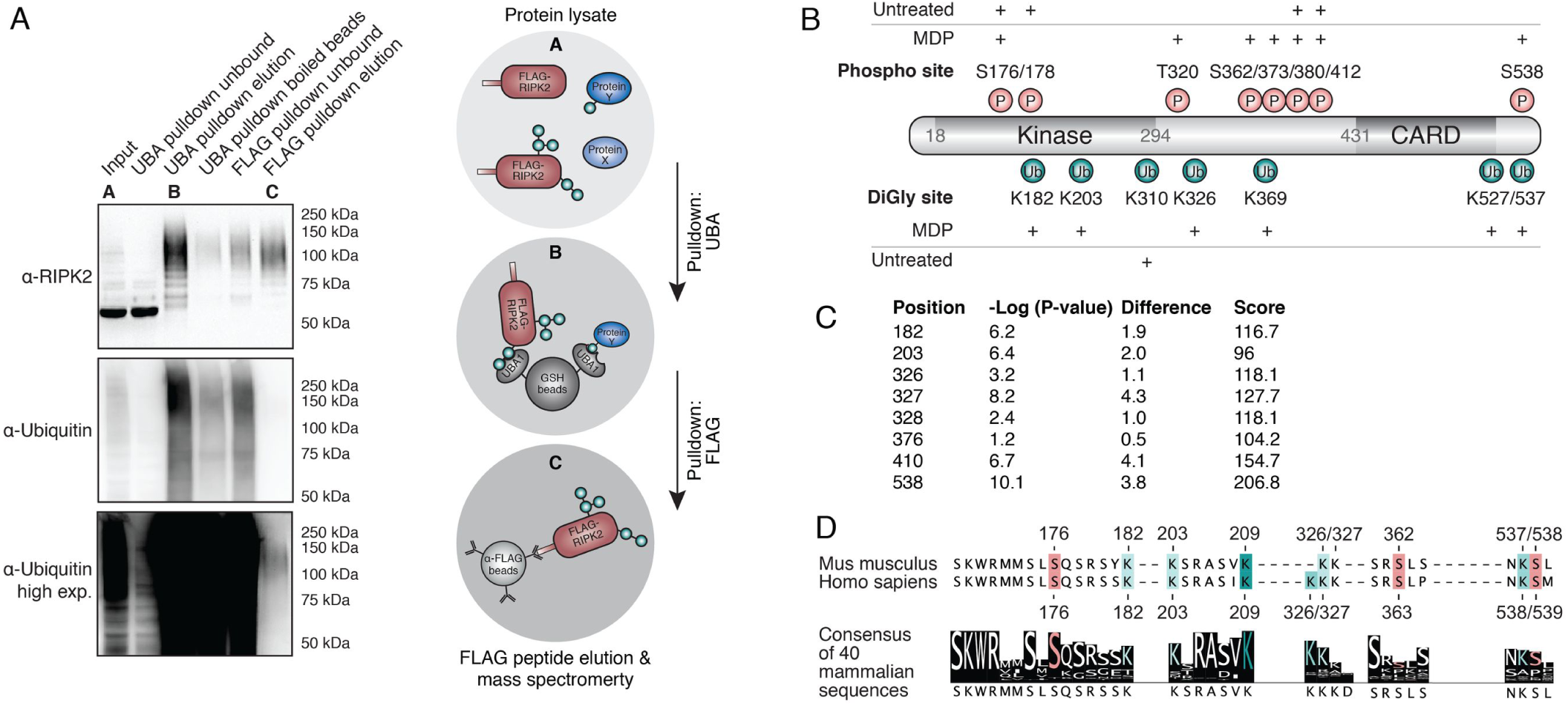
Identification of RIPK2 PTMs during NOD signaling by mass spectrometry. A Two step enrichment to isolate ubiquitinated RIPK2 from BMDMs. FLAG-RIPK2 BMDMs were sequentially subjected to ubiquitin enrichment (UBA, B) and FLAG (C) pulldown then FLAG pulldown (C) prior to protein elution and subsequent mass spectrometry analysis. B Schematic representation of RIPK2 PTMs detected in MDP-stimulated vs. unstimulated BMDMs. Red: phosphorylation, green: ubiquitination. N = 3 experiments C Table of diGly-modifications on RIPK2 determined by diGly proteomics in L18-MDP stimulated vs. unstimulated THP-1 cells. D Sequence conservation of stimulation-dependent modified serine (red) and lysine (green) residues amongst mammals. The degree of conservation is indicated by color saturation.

Tryptic digests of these samples were then generated and analysed by mass spectrometry, revealing a substantial enrichment of ubiquitin and RIPK2 peptides. To specifically determine stimulation-dependent PTMs on RIPK2, this dataset was compared to datasets obtained from FLAG pulldowns of unstimulated BMDMs. In unstimulated BMDMs, only one single K-Є-diglycine site (diGly, diglycine remnant on lysine after tryptic digestion of ubiquitinated proteins) on RIPK2 was observed at the C-terminal end of the kinase domain (K310). In contrast, MDP-stimulated BMDMs revealed multiple RIPK2 ubiquitination sites within the kinase domain (K182, K203), in the intermediate region (K326, K369) and in the CARD (K527, K537) (**Fig. 2B**). The function of most of the detected sites is still uncharacterized; only one recent study showed that human THP-1 cells expressing a K410R/K538R double mutant (human K538 corresponds to K537 in mice) displays reduced NF-*κ*B activation and cytokine production. We were not able to detect the previously described ubiquitination site K209 [27] using our stringent protocol.

Phosphor-peptides on RIPK2 were detected in both unstimulated and MDP-stimulated BMDMs. Stimulation-dependent phosphorylation was detected in the intermediate region (T320, S362, S373) and the CARD (S539). The activation loop of RIPK2 was phosphorylated on two residues (S176, S178), however, the phosphor-peptides were discovered in stimulated as well as in unstimulated cells. This was surprising since phosphorylation of the kinase activation loop is associated with RIPK2 activation [17, 35].

To further confirm the physiological importance of the RIPK2 ubiquitination sites, we examined the ubiquitome of the human monocytic cell line THP-1 employing diGly proteomics. In unstimulated cells, we did not detect any diGly sites on RIPK2 but we consistently observed several diGly marks after L18-MDP stimulation (**Fig. 2C**). The sites identified in THP-1 cells reflected our results obtained using the sequential pulldown protocol in mouse cells, further validating our initial approach. All previously detected stimulation-dependent diGly modifications on murine RIPK2 residues, that are conserved in humans, were also detected in L18-MDP stimulated THP-1 cells (K182, K203, K326, K538). Additionally, two diGly-modified lysines (K376, K410) that are not conserved in mice were found. Again, with this second protocol, we were not able to detect ubiquitination of K209.

To assess the impact of individual RIPK2 PTMs on NOD signaling we generated mutants of lysine and serine residues that we found to be modified upon NOD2 stimulation and highly conserved amongst mammals (**Fig. 2D**). Additionally, we included conserved sites that have previously been associated with RIPK2 activation (**Fig. 2D**, [17] [18] [27].). These corresponded to K182, K203, K326, K327, S363, K538, S539, and K209 in humans.

### Most PTMs on RIPK2 are redundant for NOD2 signaling

To test the function of human RIPK2 PTMs in a close-to-endogenous setting, we generated RIPK2-deficient THP-1 cells by transient transfection with Cas9 and RIPK2 sgRNA encoding plasmids and confirmed the knock-out of RIPK2 by Western blot. These knock-out cells were then transduced with doxycycline-inducible RIPK2 constructs to express RIPK2 at levels similar to wild-type THP-1. In contrast to overexpression studies, expression of RIPK2 alone did not autoactivate NF-*κ*B, but additional treatment with L18-MDP induced transient phosphorylation of p65 and I*κ*Bα and degradation of I*κ*Bα (**Fig. 3A**). Upon MDP stimulation, cells reconstituted with all mutant forms of RIPK2 activated NOD signaling normally, except the previously described K209R mutant. When tested for IL-8 secretion, all mutants, except K209R, also produced equal levels of IL-8 when compared with wild type THP-1 cells and with RIPK2-deficient cells reconstituted with wild type RIPK2 (**Fig. 3B**). Notably, RIPK2 K209R was the only form of RIPK2 which seemed to present in a second, higher molecular weight band after stimulation, similar to recently described Riposomes [36]. To test the impact of each individual modified site on the ubiquitination pattern on RIPK2 during NOD signaling, cells were subjected to UBA pulldowns and analysed by Western Blot. Stimulation with L18-MDP led to distinct RIPK2 polyubiquitination (**Fig. 3C**) and the removal of single ubiquitination or phosphorylation sites, besides K209, did not affect RIPK2 ubiquitination.

**Figure 3:**
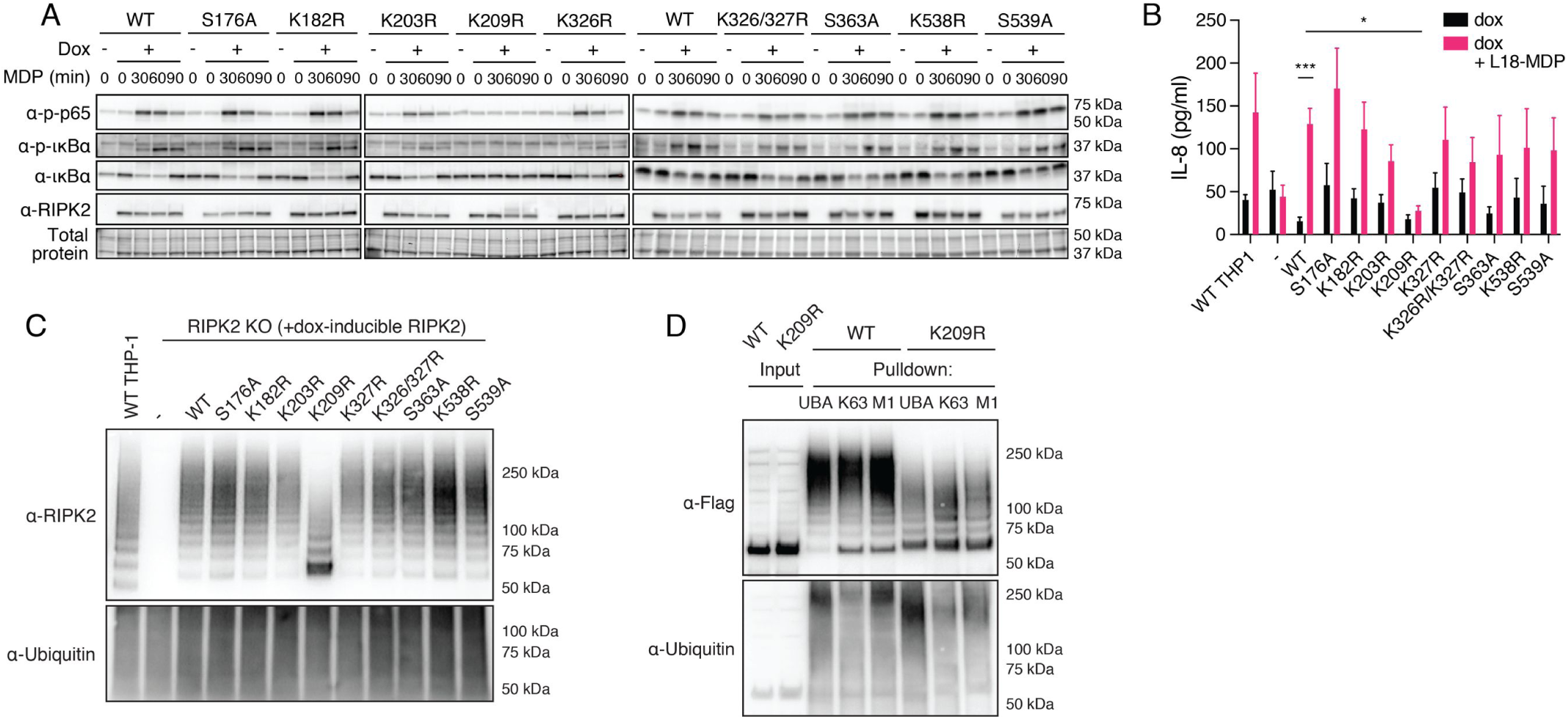
Characterization of RIPK2 diGly- and phosphosite mutations. A Activation of the NF-*κ*B pathway by RIPK2 Lysine- and phosphosite mutants. RIPK2-deficient THP-1 cells reconstituted with wild-type RIPK2 or RIPK2 mutants were stimulated with L18-MDP, harvested at indicated time points and activation of the NF-*κ*B pathway was analyzed by immunoblotting. B IL-8 production of wild-type THP-1 and RIPK2-deficient THP-1 cells reconstituted with wild-type RIPK2 or RIPK2 mutants and stimulated with L18-MDP was assessed by ELISA. N = 4-8 experiments. Shown is average ± SEM. ns = P > 0.05; * = P 0.05; ** = P 0.01; *** = P 0.001; **** = P 0.0001. RIPK2 ubiquitination determined by UBA pulldown in L18-MDP stimulated RIPK2-deficient cells reconstituted with wild-type or mutant RIPK2. Detection of K63- and M1-linked ubiquitin chains by UBA pulldown or pulldown with ubiquitin chain type-specific antibodies in L18-MDP stimulated RIPK2-deficient cells reconstituted with RIPK2 K209R.

It was previously reported that K209 is critical for RIPK2 ubiquitination and NOD signaling [27]. However, we observed that residual ubiquitination of RIPK2 was still present. Since RIPK2 K209R failed to activate NF-*κ*B and to produce cytokines and displayed reduced ubiquitination, we hypothesized that this mutation led to a loss of the critical K63- or M1-linked ubiquitin chains. To test this hypothesis, cells reconstituted with wild-type RIPK2 or K209R RIPK2 were stimulated with L18-MDP and subjected to either UBA pulldown or to pulldowns with K63- or M1-chain-specific antibodies [37, 38]. Compared to wild-type RIPK2, the ubiquitination of K209R was reduced in all pulldowns (**Fig. 3D**). However, both K63- and M1-linked chains were still present and appeared to be equally abundant on wild-type and K209R RIPK2.

### K209 and I212 form a regulatory interface critical for signal transduction

Since neither we nor others were able to detect ubiquitination of K209 by mass spectrometry [19], and mutation of K209 only reduced K63 and M1 linked ubiquitination of RIPK2 yet this same mutation abolished NF-*κ*B activation and cytokine production, we hypothesised that K209 is not a critical ubiquitination site but serves some other function. K209 is located at the N-terminal end of αE helix in the C-lobe of the kinase domain (**Fig. 4A**; PDB 4C8B; [39]). It is part of a hydrophobic pocket formed by amino acids of helix αE and amino acids in the loop between helix αE and αEF, which are suggestive of a regulatory interface. To test the hypothesis that disruption of this region prevents NOD2 signaling, we generated mutations of amino acids K209 and I208 that contribute to the make of this pocket. While K209 is invariant among vertebrates, I208 is replaced by valine in most vertebrate RIPK2 sequences (**Fig. 4B**). We also mutated I212, which is part of the αE helix with its side chain sitting deep within the pocket. I212 is highly conserved and only replaced by other hydrophobic amino acids, such as valine or methionine, in some vertebrate species.

**Figure 4:**
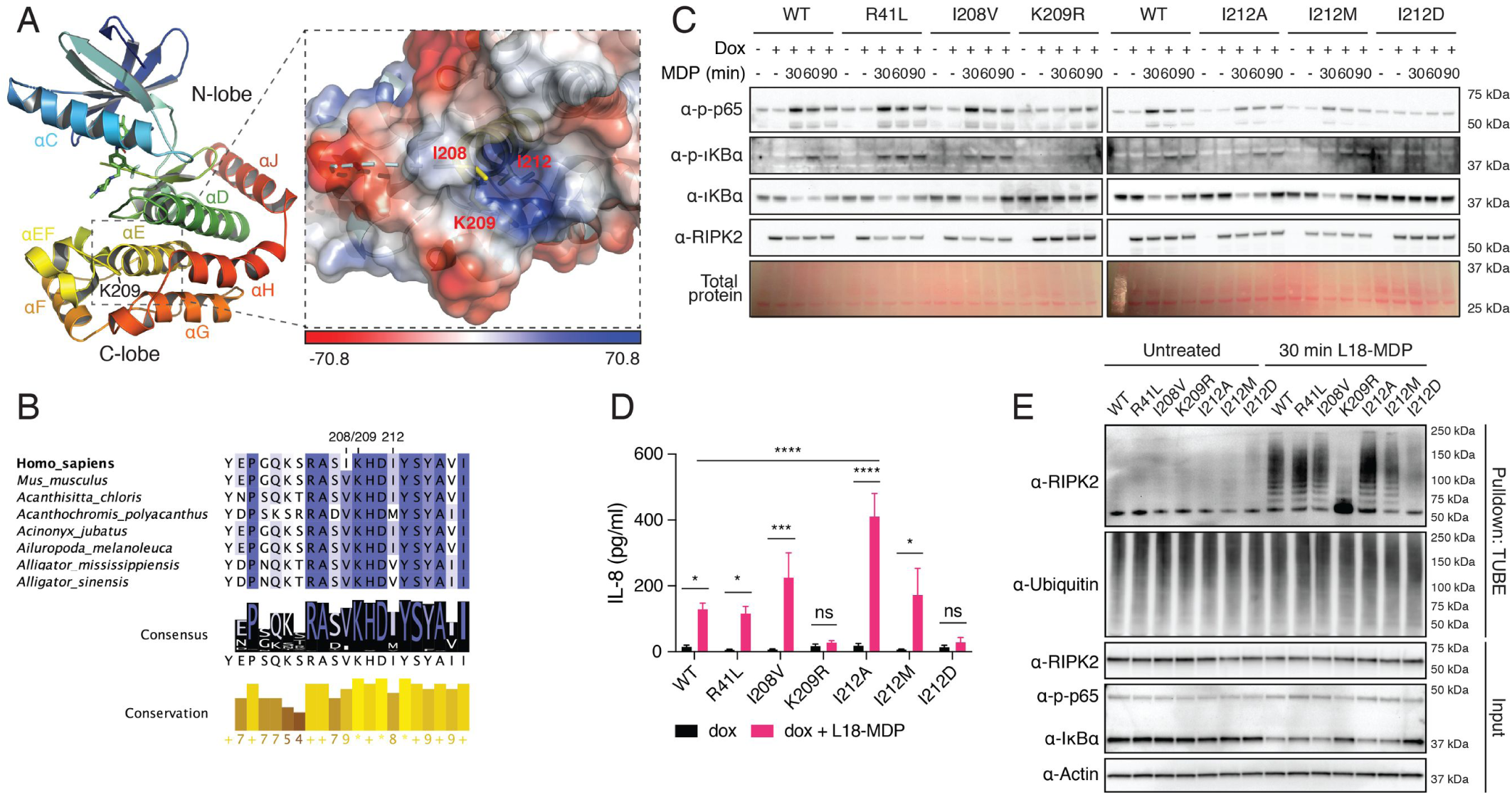
Functional studies of RIPK2 mutations introduced in close proximity to K209R. A: Structural features of the RIPK2 kinase domain (left) and location of K209 within a hydrophobic pocket between helices αEF and αE (right). Shown is chain B of the RIPK2 kinase in complex with ponatinib from PDB:4C8B. The electrostatic interaction potential is shown as a blue to red gradient. B Conservation of amino acids creating a hydrophobic pocket helices αEF and αE. Degree of conservation among 227 vertebrate species indicated by background colour saturation. C Activation of the NF-*κ*B pathway by RIPK2 pocket mutants. RIPK2-deficient THP-1 cells were reconstituted with wild-type RIPK2 or RIPK2 mutants, stimulated with L18-MDP, harvested at indicated time points and analyzed by immunoblotting. D IL-8 production of RIPK2-deficient THP-1cells reconstituted with wild-type RIPK2 or RIPK2 mutants and stimulated with L18-MDP was assessed by ELISA. N = 3-8 experiments. Shown is average ± SEM. ns = P > 0.05; * = P ≤ 0.05; ** = P ≤ 0.01; *** = P ≤ 0.001; **** = P ≤ 0.0001. E Ubiquitination of RIPK2 pocket mutants. Unstimulated or L18-MDP stimulated RIPK2-deficient THP-1 cells were reconstituted with wild-type or mutant RIPK2, left unstimulated or stimulated with L18-MDP and subjected to UBA pulldown and immunoblotting.

We reconstituted RIPK2-deficient THP-1 cells with RIPK2 mutants generated and tested them for NF-*κ*B activation, RIPK2 ubiquitination and cytokine secretion. The expression levels of these RIPK2 mutants seemed equivalent **(Fig. 4C**). Conservative I208V and I212M mutations did not prevent RIPK2 ubiquitination, activation of NF-*κ*B or the production of IL-8 after MDP stimulation. However, cells that expressed the RIPK2 I212A mutant produced significantly more IL-8 than cells expressing wild-type RIPK2 (**Fig. 4D**). Furthermore, the substitution of isoleucine 212 with an aspartic acid (I212D) completely blocked RIPK2 ubiquitination and NF-*κ*B activation. These results suggest that the C-lobe pocket may act to recruit an E3 ligase, such as XIAP.

### Mutation of K209 and I212 disrupts XIAP binding

The equivalent expression levels of the RIPK2 mutants suggested that the structural integrity of RIPK2 was not overtly compromised. However, to exclude a trivial explanation for the complete lack of signaling of the I212D mutant in particular, and more rigorously show that the mutants retained their structural integrity, we devised an intracellular thermal stability test (**Fig. 5A**) based on reports that kinase inhibitors can increase the thermal stability of their targeted kinases [40-44]. At the physiological temperature of 37°C, wild-type RIPK2 and all RIPK2 mutants were expressed at equal levels. With increased temperature, all mutants displayed similar stability compared to wild-type RIPK2. In particular, all I212 mutants, I212A (activating), I212M (residue in Damselfish) and I212D (inactivating) had an almost identical thermal stability profile, strongly suggesting that the mutations of this pocket had not caused major structural disruption.

**Figure 5:**
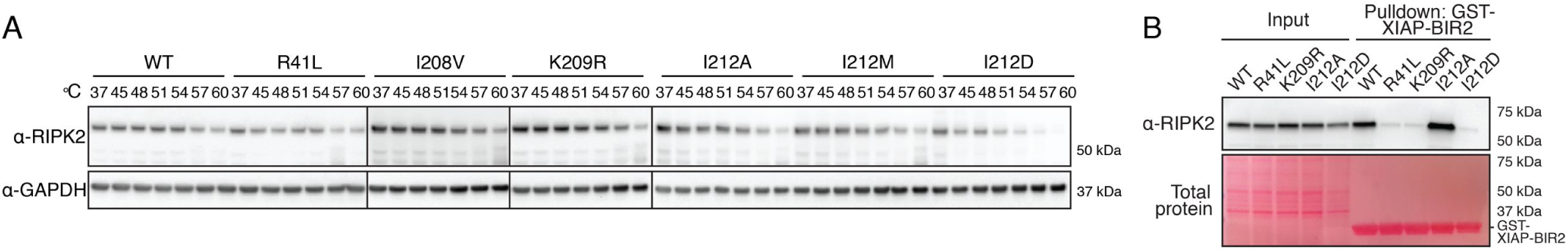
RIPK2 K209 and I212 mediate XIAP binding. A: Thermal stability assay using selected RIPK2 mutants. THP-1 cells expressing RIPK2 were subjected to heat treatment and the non-denatured fraction was analysed by immunoblotting. B: Binding of RIPK2 to XIAP-BIR2. Lysates from RIPK2-deficient THP-1 cells reconstituted with WT or mutant RIPK2 were subjected to pulldown experiments with recombinantly expressed XIAP-BIR2 and analysed by immunoblotting.

XIAP is indispensable for NOD2 responses *in vitro* and *in vivo* [22, 24, 45] and binds to the RIPK2 kinase domain via its Baculovirus IAP Repeat 2 (BIR2) domain [19, 22]. To test whether mutation of K209 and I212 disrupted XIAP binding, a purified, recombinant, GST-coupled BIR2 domain of XIAP was used to precipitate RIPK2 from THP-1 lysates (**Fig. 5B**). Wild-type RIPK2 was strongly enriched by XIAP-BIR2 pulldown, and I212A was even more abundant, correlating with the increased stimulus dependent ubiquitination of this mutant and increased cytokine secretion compared to wild-type RIPK2. Binding of K209R and I212D to the BIR2 of XIAP was, however, drastically reduced and comparable to R41L, a RIPK2 mutant previously shown to have impaired binding to XIAP [20]. This binding site for XIAP, involving R41 and R36, is in the N-lobe of the RIPK2 kinase domain and therefore quite separate (approximately 40 Å) from the C-lobe pocket, suggesting that XIAP may bind multiple sites on RIPK2. Alternatively, K209 and I212 could be important to maintain RIPK2 in a conformation that favours interactions with XIAP.

## Discussion

The molecular requirements for immune signaling through NOD1 and NOD2 have been studied in great detail. It is widely accepted that post-translational modification, particularly ubiquitination of the adaptor protein RIPK2, is the critical regulatory mechanism that determines signaling outcomes. However, while multiple ubiquitin E3 ligases and deubiquitinases (DUBs) have been suggested to regulate RIPK2 ubiquitination, the contribution of individual ubiquitination sites on RIPK2 has not been characterized in endogenous systems. Our work closes this gap and we confirm by two complementary, mass-spectrometry-based approaches in endogenous systems, that RIPK2 is ubiquitinated on multiple lysine residues. This is not unusual as many proteins become ubiquitinated on multiple sites during signaling [46, 47]. Typically, there appears to be flexibility in the lysine residues that can be ubiquitinated and often, as was the case here, mutation of a single lysine has little impact on signaling. This is believed to be because E3 ligases are not usually restricted to a specific motif, in the way that for example kinases or caspases are, and can therefore be promiscuous in the lysine that they modify [48, 49]. In line with this, we show that most single phosphorylation and ubiquitination events on RIPK2 are redundant. There are some exceptions to this general rule, for example, mutations of K377 in RIPK1 has a profound effect on RIPK1 ubiquitination and TNFR1 induced activation of NF-*κ*B [50]. It has been assumed that K209 was another similarly special residue because the K209R mutation blocked overexpression-induced NF-*κ*B activation and RIPK2 ubiquitination [27] and the mutant has a loss of function phenotype when overexpressed together with ubiquitin in HEK 293T cells [21]. However, unlike K377 in RIPK1, K209 is on the C-lobe of the kinase domain of RIPK2, which is already indicative of a different function of this site and this prompted us to closely look at the structural features of the region surrounding K209 of RIPK2. We discovered that K209 is part of a putative interaction pocket, which could explain the observed loss-of-function phenotype of the K209R RIPK2 mutant and the distinct function of K209 compared to all other ubiquitinated lysines on RIPK2.

Our data confirm previous reports that K209 is indispensable for immune signaling through NOD2, and also that RIPK2 ubiquitination is impaired when K209 is mutated to an Arginine. While this initially led us to the conclusion that K209 is directly ubiquitinated [27], we propose a different mechanism. The reason for this is twofold: Firstly, we used two multi-replicate complementary experimental approaches to determine the phosphorylation- and ubiquitination signature on RIPK2 upon activation. The fact that we identified identical diGly sites on RIPK2 in mouse (BMDMs) and in human (THP-1) gives high confidence in our datasets. Despite the high reproducibility of the 2 cell lines, in none of our experiments did we observe a diGly site corresponding to K209. Secondly, mutation of a residue in close proximity, but not directly affecting K209, resulted in an even more dramatic impact on RIPK2 ubiquitination than mutation of K209 itself. This suggests that structural integrity of this region is critical for signal transduction. To this end, we cannot exclude that our I212D mutation rendered K209 inaccessible for ubiquitination, but further studies will be required to examine this experimentally.

It must be noted that we not only failed to detect ubiquitinated K209, but we also did not detect unmodified K209 in our datasets on MDP-stimulated BMDMs or THP-1 cells. Nevertheless, when analysing other datasets that identified RIPK2 in deep proteome and pan-kinome experiments (utilising broad specificity kinase inhibitors to enrich for kinases), this region was readily identifiable with an unmodified tryptic peptide [51-54]. This could be because we specifically enriched for activated RIPK2 in our protocols and we did not use kinase inhibitors in our approaches. We therefore cannot definitively refute the idea of K209 ubiquitination, although the current evidence suggests that there may be as-yet other unidentified post-translational modifications hindering identification. Lastly, the fact that a third method did not lead to the detection of a diGly site on RIPK2 K209 [19], hints to the direction that this particular portion of RIPK2 is more complex than currently understood.

An alternative explanation for the loss of the signaling potential of the K209R and I212D mutation is instability due to misfolding or impairment of the dimerization potential. Recently cellular thermal shift assays (CETSA) have been described to monitor changes of stability of signaling proteins, in particular kinases, upon binding to small molecules [41-44]. We modified this assay to examine the effects of mutation on protein stability as a marker for structural integrity, and we think that it could be more widely used to confirm that a particular mutation does not affect structural integrity. In this case, we were able to show that the stability of the K209R or the I212D mutation is not impaired. Expression levels at physiological temperatures are equal to wild-type RIPK2, and the thermostability was not significantly changed in the mutant forms.

The E3 ligase XIAP is a critical component for NOD signaling [22, 24]. Our experiments showed that mutation of K209 or I212 impairs RIPK2 binding to the BIR2 of XIAP. These data therefore strongly suggest that both amino acids are directly or indirectly implicated in the binding of XIAP during NOD signaling. However, the structural determinants underlying the interaction between the two proteins remain unclear. The XIAP-BIR2 domain contains a deep and distinctive hydrophobic cleft that typically mediates binding to proteins harbouring an IAP-binding motif (IBM) as present in caspases or SMAC [55, 56]. However, RIPK2 does not contain such a motif. It is possible that XIAP binds to RIPK2 in a non-canonical fashion, independently of an IBM. Such a mechanism has been shown with the caspase Dronc and the XIAP homolog DIAP1 in *Drosophila melanogaster*. Binding of Dronc to DIAP1 is mediated via a short 12-residue peptide located between the CARD and the protease domain of Dronc that binds into the hydrophobic cleft of the DIAP in a similar fashion as observed in IBM-containing proteins. Therefore, an explanation could be that the RIPK2-XIAP interaction occurs in a similar, non-canonical fashion, however the region around K209 and I212 has the shape of a pocket. The binding mode of RIPK2 and XIAP-BIR2, therefore, would be completely different from the previously observed interactions. Alternatively, XIAP could bind somewhere else, and mutation of K209 and I212 might lead to conformational changes that have an impact on the structure or orientation of the interaction interface. A region in the N-lobe of RIPK2, in particular residues R36 and R41, was identified as a critical interaction region for the BIR2 of XIAP [20]. We used the reported R41L mutation as a control for our assay and confirmed the impaired binding of XIAP BIR2 to this mutant. In all reported crystal structures, the RIPK2 kinase domain is organised in head-to-tail dimers (PDB: 4C8B, 5AR4, 6ES0, 6FU5, 5J79) and the regions around R36 and R41 are quite far apart from K209. According to these structures, it is unlikely that the BIR2 of XIAP binds both areas simultaneously, but the reason for reduced binding of the K209R and I212D mutant could be due to conformational impairment.

In recent studies, higher order intracellular signalling platforms consisting of RIPK2 and NOD receptors were described [36, 57, 58]. In particular, inhibition of XIAP by either siRNA or by SMAC mimetic compounds led to RIPK2-containing speck-like structures in cells, termed Riposomes [36]. Here we described two mutations that perturbed the interaction between RIPK2 and XIAP. On close examination, RIPK2 K209R seems to accumulate in a higher order band in a Western Blot after stimulation, consistent with the hypothesis that in the absence of XIAP, RIPK2 is moved to Triton-insoluble Riposomes. However, the RIPK2 I212D or the RIPK2 R41L mutations, which reduce RIPK2-XIAP interaction to a similar level to the RIPK2 K209R mutation, did not lead to an equivalent band in Western blots after stimulation. Therefore, these new XIAP binding mutants should be useful to shed new light into the mechanism and function of Ripoptosome formation and their relevance for NOD signalling.

Taken together, our work draws a detailed map of post-translational modifications on RIPK2 during NOD signaling, and we provide evidence that most single phosphorylation and ubiquitination events on RIPK2 are redundant in systems with close to endogenous levels of RIPK2. We identified a regulatory interface on RIPK2, which dictates the crucial interaction with XIAP. This interface includes a pocket-shaped region around residues I212 and K209 on the C-lobe of RIPK2’s kinase domain. Our findings give an explanation to the conundrum why mutations of K209 reduce RIPK2 ubiquitination and block NOD signaling, yet ubiquitination of K209 has never been detected. As interfering with the RIPK2-XIAP interaction has emerged as a strategy to inhibit NOD signaling, it is tempting to speculate that this region could be targeted with small molecules for the treatment of diseases with increased NOD signalling.

## Materials and methods

### Generation of FLAG-RIPK2 CRISPR knock-in mice

The FLAG-RIPK2 mice were generated by the MAGEC laboratory (WEHI) as previously described [59] on a C57BL/6J background. To generate FLAG-RIPK2 mice, 20 ng/μl of Cas9 mRNA, 10 ng/μl of sgRNA (TGAACGGGGACGCCATCTGC;) and 40 ng/μl of oligo donor (GCCGCCCCGGGACCTAGCGCCGCGGCCAGGGTCGGGCGGAGCCGCCGCGCAGCCGGAGCCATGGACTACAAAGACGATGACGATAAAGGATCTACCAACGGGGACGCCATCTGCAGCGCGCTACCCCCCATCCCGTACCACAAGCTCGCCGACCTG) were injected into the cytoplasm of fertilized one-cell stage embryos. Twenty-four hours later, two-cell stage embryos were transferred into the oviducts of pseudo-pregnant female mice. Viable offspring were genotyped by next-generation sequencing.

### Cell culture, generation of BMDMs, and stimulation protocols

Wild type THP-1 cells and 293T cells were sourced from ATCC™. THP-1 cells were cultured in RPMI supplemented with 8% FBS and antibiotics (penicillin, streptomycin, GIBCO) at 37°C with 10% CO_2_ in a humidified incubator. 293T cells were cultured in DMEM (GIBCO) with 8% FBS in the same conditions. BMDMs were generated from the femur and tibiae of mice and cultured for six days in DMEM (InvivoGen) supplemented with 8% FBS (GIBCO) and 20% L929 supernatant and antibiotics (penicillin, streptomycin). Cells were then detached using trypsin-EDTA and re-plated in 12- and 24-well tissue culture plates. Replated cultures of BMDMs were primed with murine interferon-*γ* (5 ng/mL, R&D Systems) over night and again two hours before stimulation with MDP (10 μg/mL, InvivoGen). THP-1 cells were stimulated with L18-MDP (200 ng/mL, Bachem).

### Generation of RIPK2 CRISPR knock-out THP-1 cells

RIPK2-deficient THP-1 cells were generated using a CRISPR/Cas9 based knockout workflow as previously described [60]. Briefly, a sgRNA (GACCTGCGCTACCTGAGCCGCGG) targeting RIPK2 was designed. THP-1 cells were nucleofected with one plasmid expressing sgRNA and one expressing mCherry-Cas9 (pLK0.1-gRNA-CMV-GFP, CMV-mCherry-Cas9) using the SG Cell Line 4D-Nucleofector™ X Kit S and a 4D-Nucleofector X unit. mCherry positive cells were sorted and cloned by limiting dilution. After identifying clones, cells were replated and grown to identify RIPK2 knock-out clones by assessing RIPK2 expression on Western Blot.

### Intraperitoneal MDP injections

All in vivo experiments were performed according to the guidelines of the animal ethics committee of WEHI, ethics approval (2011.014, 2014.004 and 2017.004). Sex and age matched littermate controls were used within each experiment. For in vivo MDP challenge: wild-type, RIPK2 knock-out and FLAG-RIPK2 knock-in (hetero- and homozygous) mice were administered MDP (5 mg/kg, i.p. in 200 μL PBS, Bachem) or PBS and sacrificed after 4 hours. Peripheral blood was collected by cardiac puncture.

### Western blotting

Following stimulation, cells were lysed in 2 x SDS lysis buffer (126 mM Tris-HCl pH 8, 20% v/v glycerol, 4% w/v SDS, 0.02% w/v Bromophenolblue, 5% v/v 2-mercaptoethanol) and subjected to repeated freeze/boil cycles. Samples were separated using SDS PAGE and transferred to polyvinylidene fluoride (PVDF) membranes. The following antibodies were used for probing: rabbit anti RIPK2 (4142S, Cell Signaling Technology), rabbit anti RIPK2 (SC 22763, Santa Cruz), mouse anti FLAG (F1804, Sigma), anti β actin (A-1978, Sigma), rabbit anti p65 (631213, Upstate), rabbit anti phospho p65 (3033, Cell Signaling Technology), rabbit anti phospho p38 (9211, Cell Signaling Technology), mouse anti phospho IκBα (9246, Cell Signaling Technology), rabbit anti IκBα (9242, Cell Signaling Technology), mouse anti ubiquitin (3936, Cell Signaling Technology), rabbit anti GAPDH (2118, Cell Signaling Technology), human anti-K63-linked ubiquitin (Apu3.A8, Genentech), human anti M1-linked ubiquitin (1F11, Genentech), anti K27-linked ubiquitin (ab18153, abcam), goat anti mouse Ig (1010-05), goat anti rabbit Ig (4010-05) and goat anti rat Ig HRP (horseradish peroxidase, 3010-05, Southern Biotech), goat anti human Ig (109-035-003, Jackson Immunoresearch).

### Cytokine measurement by ELISA

Cytokines from mouse serum or cell culture supernatant were measured by ELISA kits for IL-6, IL-8, TNF and MCP-1 respectively (Invitrogen) according to the manufacturer’s instructions. Sera and supernatants were diluted 1:10 for MCP-1 measurements.

### Immunoprecipitation of RIPK2 from mouse tissues

Organs from 6 weeks old wild-type C57BL/6 mice and FLAG-RIPK2 knock-in mice were lysed in IP buffer (150 mM NaCl, 50 mM Tris pH 7.5, 10% glycerol, 1% Triton X-100, 2 mM EDTA, all from SIGMA) supplemented with protease inhibitors (cOmplete protease inhibitor cocktail, Roche) using a TissueLyser II (Qiagen). Samples were clarified by centrifugation at 17000 x g for 30 min, the protein concentration was assessed using a BCA assay (Thermo Fisher) and 2 mg of protein per lysate were subjected to anti-FLAG immunoprecipitation using 15 ul of packed magnetic anti-FLAG beads (M2, Sigma) for 4h. Beads were washed 3 times in IP buffer, eluted with 2x SDS sample buffer and subjected to immunoblotting.

### Ubiquitin enrichments (UBA, TUBE and pulldowns with K63- and M1-specific ubiquitin antibodies)

20*10^6^ THP-1 cells were treated with doxycycline (200 ng/ml) for 5 h and stimulated with L18-MDP (200 ng/ml, Invitrogen) for 30 min, washed in PBS and lysed in 1-2 ml IP buffer (150 mM NaCl, 50 mM Tris pH 7.5, 10% glycerol, 1% Triton X-100, 2 mM EDTA) with protease and phosphatase inhibitors and 5 mM n-ethylmaleimide (NEM). Samples were clarified by centrifugation at 17000 x g for 15 min and added directly to 20 ul packed glutathione sepharose beads pre-bound with 100 ug GST-TUBE or GST-UBA. Beads were incubated on a rotating wheel at 4°C for at least 2 h, washed 3 times with IP buffer and eluted with 2x SDS sample buffer.

To enrich for K63- and M1-linked ubiquitin species, 20*10^6^ THP-1 cells were treated with doxycycline (200 ng/ml) for 5 h and stimulated with L18-MDP (200 ng/ml, Invitrogen) for 30 min, washed with PBS and lysed in 1-2 ml IP buffer, as above, supplemented with 6M Urea (for anti-K63-linked ubiquitin pulldowns) or 8M Urea (for anti M1-linked ubiquitin pulldowns). Samples were clarified by centrifugation at 17000 x g for 15 min and 4 ug of anti-K63 or anti-M1-linked ubiquitin antibodies (Genentech; [37, 38]) were added, followed by incubation on a rotating wheel at 4°C for at least 2 h. Antibodies were precipitated with 10 ul of equilibrated protein G agarose (Thermo), washed 3 times in IP buffer without Urea and eluted with 2x SDS sample buffer.

### Two step enrichment of modified RIPK2

12 dishes of confluent FLAG-RIPK2 BMDMs (equivalent of approximately 25*10^7^ cells or 25 mg of total protein) were primed with IFNγ (5 ng/ml) overnight and fresh IFNγ was added the next morning for another 2 h before stimulation with MDP (10 ug/ml) for 30 min. Cells were harvested, washed in PBS and lysed in 2 ml IP buffer per dish (150 mM NaCl, 50 mM Tris pH 7.5, 10% glycerol, 1% Triton X-100, 2 mM EDTA; supplemented with protease and phosphatase inhibitors and 5 mM n-ethylmaleimide (NEM)). Lysates were clarified by centrifugation at 17000 x g for 15 min and supernatants were directly added to 100 ul packed glutathione sepharose beads pre-bound with 1 mg GST-UBA. Beads were incubated on a rotating wheel at 4°C overnight, washed 3 times with IP buffer and eluted twice with two volumes of IP buffer supplemented with 20 mM reduced glutathione (pHed to 7.5). Elutions were combined, diluted with an equal volume of IP buffer and added to 50 ul of packed magnetic anti-FLAG beads (M2, Sigma) and incubated on a rotating wheel at 4°C for 4 hours. Beads were washed 3 times with IP buffer and eluted twice with two volumes of 3x-FLAG peptide (1 mg/ml) in TBS pH 7.5.

### THP-1 diGly proteomics

Frozen pellets from 50 × 10^6^ WT THP-1 cells were lysed in 1% SDC, 100mM Tris/HCL pH 8.5, immediately boiled for 5 min at 95°C and sonication for 30s (Branson Sonifierer). Protein concentrations were estimated by tryptophan assay. For protein reduction and alkylation samples were incubated for 5 min at 45°C after addition of TCEP and CAA to a final concentration of 10 mM and 40 mM, respectively. Samples were digested using Trypsin (1:20 w/w, Sigma-Aldrich) in combination with LysC (1/100 w/w, Wako) at 37°C overnight. Protease activity was quenched by addition of four-sample volumes 1% TFA in Isopropanol. Quenched samples were loaded onto SDB-RPS cartridges (Strata™-X-C, 30mg/3mL, Phenomenex Inc), pre-equilibrated with 4ml 30% Methanol (MeOH)/1% TFA and washed with 4ml 0.2% TFA. After 2 washes with 4 ml 1%TFA in Isopropanol and 1 wash with 0.2% TFA/2%ACN samples were eluted twice with 2ml 1.25% Ammonium hydroxide(NH_4_OH)/80% ACN. Eluted samples were diluted with ddH_2_0 to a final ACN concentration of 35%, snap frozen and dried by lyophilization.

Lyophilized peptides were reconstituted in IAP buffer (50mM MOPS, pH 7.2, 10mM Na2HPO4, 50mM NaCl) and the peptide concentration was estimated by tryptophan assay. For proteome analysis, 10µg of peptide material was taken and desalted on SDB-RPS StageTips (Empore) [61]. Peptides were diluted to a final volume of 200µl with 0.2% TFA and loaded onto StageTips, followed by a wash with 200µl of 0.2% TFA and 200µl of 0.2%TFA/2%ACN, respectively. Captured peptides were eluted with 60 µl of 1.25% Ammonium hydroxide(NH_4_OH)/80% ACN and dried using a SpeedVac centrifuge (Eppendorf, Concentrator plus). Dried peptides were resuspended in buffer A* (2 % ACN/0.1 % TFA).

K-ε-GG remnant containing peptides were enriched using the PTMScan® Ubiquitin Remnant Motif (K-Є-GG) Kit (Cell Signaling Technology). Crosslinking of antibodies to beads and subsequent immunopurification was performed with slight modifications as previously described [62]. Briefly, cross-linked beads were split equally into 8 tubes (∼31µg of antibody per tube), gently mixed with 1mg peptide material (1mg/ml) and incubated for 1h at 4°C. Beads were washed twice with cold IAP and 5 times with cold ddH_2_O and peptides were eluted twice with 50µl 0.15% TFA. Eluted peptides were desalted and dried as described above and resuspended in 5µl buffer A* for LC/MS-MS analysis.

### Mass spectrometry analysis

For the two-step protocol in BMDMs, eluted protein material from pulldowns of FLAG-RIPK2 expressing BMDMs was subjected to tryptic digestion using the FASP method as previously described[63]. Peptides were lyophilised using CentriVap (Labconco) prior to reconstituting in 80 μl 0.1% FA/2% acetonitrile (ACN). Peptide mixtures were analysed by nanoflow reversed-phase liquid chromatography tandem mass spectrometry (LC-MS/MS) on an M-Class HPLC (Waters) coupled to a Q-Exactive Orbitrap mass spectrometer (Thermo Fisher). Peptide mixtures were loaded in buffer A (0.1% formic acid, 2% acetonitrile, Milli-Q water), and separated by reverse-phase chromatography using C_18_ fused silica column (packed emitter, I.D. 75 μm, O.D. 360 μm x 25 cm length, IonOpticks, Australia) using flow rates and data-dependent methods as previously described [64, 65].

For the THP-1 diGly-enriched analysis, samples were loaded onto a 50 cm reversed phase column (75 μm inner diameter, packed in house with ReproSil-Pur C18-AQ 1.9 μm resin [Dr. Maisch GmbH]). The column temperature was maintained at 60°C using a homemade column oven. Peptides were separated with a binary buffer system of buffer A (0.1% formic acid (FA)) and buffer B (80% acetonitrile plus 0.1% FA), at a flow rate of 300 nl/min. Nano flow Liquid chromatography was performed with an EASY-nLC 1200 system (Thermo Fisher Scientific), which was directly coupled online with the mass spectrometer (Q Exactive HF-X, Thermo Fisher Scientific) via a nano-electrospray source. For proteome measurements 500ng were loaded and eluted with a gradient starting at 5% buffer B and stepwise increased to 30% in 95min, 60% in 5 min and 95% in 5min. The mass spectrometer was operated in Top15 data-dependent mode (DDA) with a full scan range of 300-1650 m/z at 60,000 resolution with an automatic gain control (AGC) target of 3e6 and a maximum fill time of 20ms. Precursor ions were isolated with a width of 1.4 m/z and fragmented by higher-energy collisional dissociation (HCD) (NCE 27%). Fragment scans were performed at a resolution of 15,000, an AGC of 1e5 and a maximum injection time of 28ms. Dynamic exclusion was enabled and set to 30s.

For K-Є-GG peptide samples 2µl were loaded and eluted with a gradient starting at 3% buffer B and stepwise increased to 7% in 6 min, 20% in 49 min, 36% in 39 min, 45% in 10min and 95% in 4 min. The mass spectrometer was operated in Top12 data-dependent mode (DDA) with a full scan range of 250-1350 m/z at 60,000 resolution with an automatic gain control (AGC) target of 3e6 and a maximum fill time of 20ms. Precursor ions were isolated with a width of 1.4 m/z and fragmented by higher-energy collisional dissociation (HCD) (NCE 28%). Fragment scans were performed at a resolution of 30,000, an AGC of 1e5 and a maximum injection time of 110ms. Dynamic exclusion was enabled and set to 15s.

### MS data processing

For BMDM data sets, raw files consisting of high-resolution MS/MS spectra were processed with MaxQuant (version 1.5.8.3) for feature detection and protein identification using the Andromeda search engine [66]. Extracted peak lists were searched against the UniProtKB/Swiss-Prot *Mus musculus* database (October 2016) and a separate reverse decoy database to empirically assess the false discovery rate (FDR) using strict trypsin specificity allowing up to 2 missed cleavages. The minimum required peptide length was set to 7 amino acids. The mass tolerance for precursor ions and fragment ions were 20 ppm and 0.5 Da, respectively. The search included variable modifications of oxidation (methionine), amino-terminal acetylation, carbamidomethyl (cysteine), GlyGly or ubiquitination (lysine), phosphorylation (serine, threonine or tyrosine) and N-ethylmaleimide (cysteine). Raw MS data were also searched with PEAKS, version 8 (Bioinformatics Solutions) using a Swiss-Prot Human database and the same variable and fixed modifications as described above. A 0.1% and 1% FDR cut-off were applied at the PSM and peptide/protein levels, respectively.

Raw MS data from THP-1 cells were searched against the UniProt Human FASTA (21051 sequences) using MaxQuant (version 1.6.2.10) with a 1% FDR at peptide and protein levels. The match and alignment time window for the match between run (MBR) algorithm were set to 0.7 min and 20min, respectively. A ratio count of 2 was used for the MaxLFQ algorithm. Cysteine carbamidomethylation was defined as fixed, protein N-terminal acetylation and methionine oxidation as variable modification. In case of K-Є-GG samples, “GlyGly (K)” was additionally selected as variable modifications. Enzyme specificity was set to trypsin and two missed cleavages were allowed, while permitting a maximum of 5 modifications per peptide.

The mass spectrometry proteomics data have been deposited to the ProteomeXchange Consortium via the PRIDE partner repository [67] with the dataset identifier PXD017741.

### Recombinant protein purification

pGEX-6 P-1 or pGEX-6 P-3 plasmids encoding XIAP-BIR2, Ubiquillin-UBA1x (UBA) or Ubiquillin-UBA4x (TUBE; [68]) were transformed into BL21 (DE3þ) bacteria and grown in Super broth overnight at 37°C. Overnight culture was diluted 1:10 and grown until OD595 was 0.8. Isopropylthiogalactoside (IPTG) (0.3 mM) was added for 4 h at 30°C. Cells were pelleted and resuspended in Buffer A (50 mM Tris pH 8.0, 50 mM NaCl, 1 mM EDTA, 1 mM DTT, 10% glycerol, all from SIGMA) and sonicated. After centrifugation at 21,000gfor 30 min, the supernatant was incubated with glutathione sepharose4B (GE Healthcare) for 4 h, washed five times with Buffer A, and eluted 2×45 min with 10 mM reduced glutathione in Buffer A at 4°C.

For generation of recombinant FLAG-RIPK2 protein from HEK 293T cells, 15 cm plates of cells were transfected with 10 ug DNA (pcDNA 3.1 plasmids encoding wild-type FLAG-RIPK2 or K209R FLAG-RIPK2) using Lipofectamine 2000 (Invitrogen) at a 1:1 ratio of reagent:DNA. After 24h cells were harvested and lysed in 1-2 ml IP buffer (150 mM NaCl, 50 mM Tris pH 7.5, 10% glycerol, 1% Triton X-100, 2 mM EDTA) with protease and phosphatase inhibitors (5 mM β-Glycerophosphate, 1 mM Sodium molybdate, 2 mM Sodium pyrophosphate, 10 mM Sodium fluoride). Samples were clarified by ultracentrifugation and supernatants were supplemented with NaCl up to a final concentration of 500 mM and subjected to anti-FLAG immunoprecipitation using 100 ul of packed beads magnetic beads (anti-FLAG, M2, Sigma). After 16h beads were washed 5 times with IP buffer and eluted twice with 250 ul of 1 mg/ml 3x-FLAG peptide (Bachem) in TBS pH 7.5 and eluates were pooled.

### Generation of doxycycline-inducible cell lines

Sequences of full length hsRIPK2 with an N-terminal 3x-Flag tag were synthesized by Genscript and cloned into doxycycline-inducible lentiviral expression vectors (pF_TRE3G_rtTAAd_puro (Takara Bio)). For lentiviral transfections, 2.5ug of the plasmid of choice was transfected into HEK293T cells together with 1 ug pVSV-G and 1.5 ug pCMVΔR8.2 using an Effectene transfection kit (Qiagen). Twenty-four hours after transfection, the media was changed and virus was harvested after another twenty-four hours. Media was filtered and supplemented with polybrene (4 μg/ml). Viral media was then applied to cell lines, centrifuged for 45 minutes at 2000 rpm @ 30°C. After two days of incubation, cells were selected using 2.5 μg/ml puromycine (Sigma).

### Thermal shift assay

5*10^6^ THP-1 cells were treated with doxycycline (200 ng/ml, Sigma) for 5 h, washed in PBS and resuspended in PBS supplemented with protease inhibitors. Cell suspensions were transferred into PCR tubes and incubated for 3 minutes at a temperature gradient (37 – 60°C) in a PCR machine. Samples were cooled to room temperature, lysed by repeated freeze-thawing and centrifuged at 20000g for 20 minutes @ 4C. Supernatants were harvested and 2x SDS sample buffer was added, before analysis by Western Blot.

### XIAP-BIR2 binding assay

20*10^6^ THP-1 cells were treated with doxycycline (200 ng/ml, Sigma) for 5 h, washed in PBS and lysed in 1-2 ml IP buffer (150 mM NaCl, 50 mM Tris pH 7.5, 10% glycerol, 1% Triton X-100, 2 mM EDTA) with cOmplete protease inhibitor cocktail, Roche and phosphatase inhibitors (5 mM β-Glycerophosphate, 1 mM Sodium molybdate, 2 mM Sodium pyrophosphate, 10 mM Sodium fluoride) and 5 mM n-ethylmaleimide (NEM, Sigma). Lysates were clarified by centrifugation at 17000 x g for 15 min and added directly to 20 ul packed glutathione sepharose beads pre-bound with 100 ug GST-XIAP-BIR2. Beads were incubated on a rotating wheel at 4°C for at least 2 h, washed 3 times with IP buffer and eluted with 2x SDS sample buffer.

## Acknowledgements

We thank Prof. C. Day for the GST-BIR2 constructs and C. Gatt, K.McKenzie, T. Ballinger and C.Epifanio for assistance with animal work and the MAGEC facility of WEHI for the generation of affinity-tagged RIPK2 mice.

## Author contributions

JS, AW and UN developed the concept for this work. VH, LD, CS, FH, EC, AB and IL performed experimental work. AW oversaw the mass spectrometry work and IL the biophysical characterisation of generated RIPK2 mutants. VH, JS and UN wrote and revised the manuscript.

